# MEAHNE: MiRNA-disease association prediction based on semantic information in heterogeneous networks

**DOI:** 10.1101/2022.05.11.491444

**Authors:** Chen Huang, Keliang Cen, Yang Zhang, Bo Liu, Yadong Wang, Junyi Li

## Abstract

Prior studies have suggested close associations between miRNAs and diseases. Correct prediction of potential miRNA-disease pairs by computational methods is able to greatly accelerate the experimental process in biomedical research. However, many methods cannot effectively learn the complex information in the multi-source data, and limits the performance of the prediction model. A heterogeneous network prediction model MEAHNE is proposed to make full use of the complex information in multi-source data. We first constructed a heterogeneous network using miRNA-disease associations, miRNA-gene associations, disease-gene associations, and gene-gene associations. Because the rich semantic information in the heterogeneous network contains a lot of relational information of the network. To mine the relational information in heterogeneous network, we use neural networks to extract semantic information in metapath instances. We encode the obtained semantic information into weights using the attention mechanism, and use the weights to aggregate nodes in the network. At the same time, we also aggregate the semantic information in the metapath instances into the nodes associated with the instances, which can make the node embedding have excellent ability to represent the network. MEAHNE optimizes parameters through end-to-end training. MEAHNE is compared with other state-of-the-art heterogeneous graph neural network methods. The values of area under precision-recall curve and receiver operating characteristic curve show the superiority of MEAHNE. Additionally, MEAHNE predicted 50 miRNAs for lung cancer and esophageal cancer each and verified 49 miRNAs associated with lung cancer and 44 miRNAs associated with esophageal cancer by consulting relevant databases. MEAHNE has good performance and interpretability by experimental verification.

## INTRODUCTION

MiRNA is a type of noncoding RNA that plays an important role in the regulation of gene expression in eukaryotes [1-3]. Through the continuous improvement of biological experiment technology, the important roles of miRNAs in the occurrence and development of diseases have been revealed [4-6]. During the development of diseases, a miRNA can inhibit or promote disease by interacting with miRNA target [7-8]. Identifying the miRNAs related to a disease is of great help for prevention and diagnosis. Therefore, researchers have carried out a large number of experiments between miRNA and disease However, the number of elements in the existing miRNA set is much larger than the number of miRNAs associated with diseases, which brings great challenges to biological experiments.

Computational methods can help researchers find miRNAs that are highly likely to be associated with disease. Computational methods have gradually begun to be applied to the study of the association between miRNA and disease [9-10]. Finding high-probability miRNA-disease pairs using computational methods and then conducting biological experiments to experimentally verify these pairs can greatly increase the hit rate of biological experiments and save costs. The computational research methods for predicting the association between miRNA and disease are mainly divided into three categories: prediction based on similarity measures, the relation-based representation learning method, and graph neural network method. Our method processed data from different sources into heterogeneous network and uses heterogeneous graph neural network methods to learn the representation of the network.

Based on similarity measures method, the central idea of the method is miRNA with similar functions may be associated with the same disease. Jiang et al. [11] established a miRNA functional similarity matrix and a miRNA-disease adjacency matrix to form a network, and calculated the similarity score in the network. Chen et al. [12] designed a prediction model, which integrates miRNA functional similarity, disease semantic similarity, and Gaussian interaction profile kernel similarity between disease and miRNA. Then calculate within-score and between-score between miRNA and disease to make prediction.

The relationship-based representation learning method is used for learning the relationship structure between miRNAs and disease. The relationship between miRNAs and disease can be modeled as a relationship network or as a relationship matrix. During representation learning of the relationship network, the relationship between miRNAs and disease is first integrated into a network. Chen et al. [13] regarded disease-related miRNAs as seeds, and used these seeds as starting points to perform a restarting random walk on the miRNA functional similarity matrix. You et al. [14]in order to alleviate the problem of sparse connections in the similarity network, use Gaussian profile kernel similarity to supplement the functional similarity matrix of miRNA and disease. And based on the obtained similarity matrix and miRNA-disease adjacency matrix, a heterogeneous network is established. They used the Depth-First-Search method to predict potential miRNA-disease associations as they traveled through the network. Another very popular approach is representation learning of the relationship matrix. This method builds the miRNA disease relationship into a matrix and decomposes the matrix. Wu et al. [15] completes and optimizes the miRNA-disease adjacency matrix. They use the miRNA functional similarity matrix and the disease semantic similarity matrix and the KNN method to complete the miRNA-disease adjacency matrix. And they used the collaborative matrix decomposition method to obtain the representation matrix of miRNA and disease. Chen et al. [16] combined miRNA functional similarity, disease semantic similarity, and Gaussian interaction profile kernel similarity calculations as the comprehensive similarity of miRNA and disease, add the similarity into the miRNA-disease adjacency matrix, and finally decompose the adjacency matrix

In the graph neural network learning method, the miRNAs and disease are first built into a graph, then the graphs are learned using graph neural network (GNN) methods [17-20]. The node representation of the GNN method fuses the structural information and the attribute information of the network. At the same time, the graph neural network end-to-end training method can also be used to optimize all the parameters in the model. Therefore, the learning ability of graph neural networks is very powerful. Li et al. [21] established a miRNA functional similarity matrix and disease semantic similarity matrix into a graph, and used GCN [17] to learn the structure information of the graph; they then used the structure information as the input for a multi-layer neural network to obtain a low-dimensional representation of miRNA and disease. To effectively integrate heterogenous miRNA and disease information, Li et al. [22] designed a graph encoder, which contains an aggregator function and a multi-layer perceptron that aggregates node neighborhood information to generate a low-dimensional embedding of miRNAs and diseases.

Many methods learn on homogeneous data, and isomorphic graph neural networks cannot adapt well to the complex associations of heterogeneous networks obtained when using multi-source heterogeneous data. To learn the semantic information generated by the complex associations in the network, the heterogeneous neural network performs multi-modal information mining on the heterogeneous network by setting the metapath. Each metapath represents a semantic type. Multiple subgraphs are sampled from the heterogeneous network according to the set of multiple metapaths, and then the graph neural network method is used to learn a low-dimensional representation of the nodes on the subgraphs. The concept of metapath was first proposed by Metapath2vec Dong Y et al. [23]. Metapath2vec samples multiple sequences composed of nodes from heterogeneous networks through the metapath setting, and a word representation learning model processes the sequences into low-dimensional vector representations. HAN [24], a representative heterogeneous graph neural network, processes the heterogeneous network into multiple sub-netwoks through metapath, and processes each subgraph into a graph composed of corresponding nodes of the same type. GAT [18] is then used to learn low-dimensional representations of isomorphic subgraphs, and semantic level attention is used to integrate the representations under multiple metapaths. HAN learns the semantic information in the network, and it can better represent the nodes in the network than the isomorphic neural network. However, this method processes the sub-graphs under the metapath into isomorphic graphs, ignoring all intermediate nodes, resulting in a large amount of information being ignored. This problem is also called the early-summarization problem [25]. MAGNN [26] is a heterogeneous neural network model based on HAN. To solve the problem of missing information in the intermediate nodes on the metapath subgraph, MAGNN rotates the intermediate nodes of each metapath instance of the subgraph. The low-dimensional embedding obtained by the rotation is regarded as the semantic information of the instance, and the semantic information is aggregated into the target nodes. In the MAGNN method, the information of all types of nodes is fused together, which leads to the loss of discrimination between the representations of different types of nodes.

The traditional heterogeneous graph neural network aggregates nodes in the network indiscriminately, which wastes the semantic information in the network. In fact, semantic information in the network can help the network to aggregate nodes more efficiently. To overcome this problem, we propose a semantic-based attention aggregation heterogeneous graph neural network to predict miRNA-disease potential association. Our main contributions are as follows:

- To fully utilize the semantic information in heterogeneous graph neural networks, we propose a semantic-based attention mechanism that utilizes the extracted semantic information to efficiently aggregate nodes in heterogeneous networks.
- In addition to aggregating neighbor node information, the semantic information extracted from metapath instances is also aggregated into nodes associated with instances. This enables nodes to have rich semantic information and adequately express relationships in heterogeneous networks.
- We design a semantic-based heterogeneous graph neural network model. By utilizing the semantic information of multiple metapaths and the semantic information of metapath instances, the relationships in the mirna-disease-gene network are fully mined. Our model can be used for the mining of large-scale multi-source biological data.

## EXPERIMENTS

In this section, we introduce several representative models of heterogeneity graph representation, compare and analyze them with our model in detail. We compare our method with other heterogeneous network embedding methods under two metrics, area under the receiver operating characteristic curve (AUC), and area under the precision-recall curve (AP), under fair conditions. And draw the Receiver Operating Characteristic(ROC) and Precision Recall(P-R) curves. Then the advantages of our model analyzed for miRNA-disease link prediction tasks in large-scale heterogeneous networks by observing and comparing experimental performances. The models we used for comparison are as follows:

Metapath2vec [23]: A structural learning method for heterogeneous networks. The network is sampled according to the set metapath to obtain a sentence composed of nodes in the network, and the sentence is used as an input for the skipgram model to obtain the final node embedding. We experimented with multiple metapaths and obtained the best performance under the metapath(mirna-disease-gene-mirna).

GAT [18]: A type of isomorphic graph neural network. This model uses the attention mechanism to assign weights to the neighbors of nodes in the spatial domain. According to the calculated weights, the neighbors in the spatial domain are aggregated. GAT uses multi-head attention, which is used to comprehensively learn the network and generate the final node representation.

HAN [24]: A heterogeneous graph neural network model that uses multiple metapaths to mine the network, separates the corresponding subgraphs, and processes the subgraphs into isomorphic graphs; GAT is then used to learn the processed graph to obtain the node representation under a single metapath, and then the attention mechanism is used to fuse the node representations under multiple metapaths.

MAGNN [26]: A heterogeneous graph neural network model. This model first uses multiple metapaths to sample the network to obtain multiple subgraphs under different source paths. To preserve the instances of each subgraph, the semantic information of each instance is rotated, and different types of rotations are rotated into the same space as the semantic information of each instance. The attention mechanism is used to aggregate the semantic information of the instances into the nodes. The problem of premature integration is alleviated. Finally, semantic level attention is used to fuse the node representations under multiple source paths.

HeCo [32]: A self-supervised heterogeneous graph neural network. The node representation of the heterogeneous network is learned from two perspectives, namely the network architecture perspective and the metapath perspective, which fully capture the information in the heterogeneous network. By using collaborative contrastive learning for node embedding from the two perspectives, network perspective and meta-path perspective are collaboratively supervised as two views. As training progresses, these two views will guide each other and co-optimize.GAEMDA [33]: An autoencoder model used on bipartite graphs. The model first projects the two types of nodes in the bipartite graph into the same space through the node transformation matrix, and aggregates the features of other types of nodes into the original embedding of the node through the encoder of the graph neural network. Finally, the prediction of potential links between nodes is done using a bilinear decoder.

The parameter settings used for the models were as follows:

The window size of the Metapath2vec model was set to 5 and the walk length to 100; each node performed 10 walks, and the number of negative samples was 5. In the GAT model, the hidden layer dimension was set to 64, the multi-head attention to 2, and the learning rate to 0.0001. HAN, MAGNN, and our method MEAHEN are heterogeneous neural networks, which are methods for metapath segmentation of the original heterogeneous network and learning of segmentation subgraphs, so we used the same parameters for all three models. Since both MAGNN and our model MEAHNE set a limit on the number of nodes in the node sampling stage, for fairness, we set the same limit for the HAN model: a maximum of 100 neighbors for each node. The node dimensions of the three models were all set to 64, with a learning rate of 0.005 and an L2 penalty weight of 0.001. In the GAEMDA model, the number of node aggregation layers was set to 2, the learning rate to 0.001, and the weight decay to 0.001. In the HeCo model, the learning rate was set to 0.001, the number of neighbor samples in the network architecture perspective to 10, and the dimension of the hidden layer to 64. Table 1 shows the experimental results of all models according to two indicators, AUC and AP, and Fig 1 shows the ROC curve and P-R curves for several linked prediction tasks.

**Table 1.**
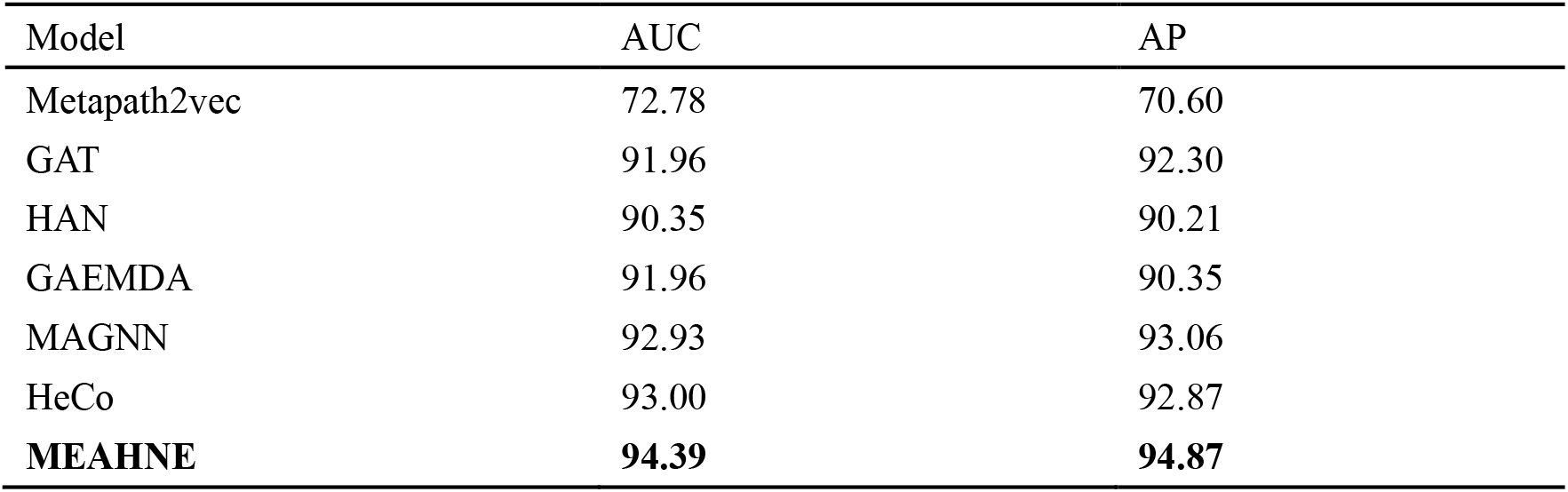
Model evaluation

**Fig 1.**
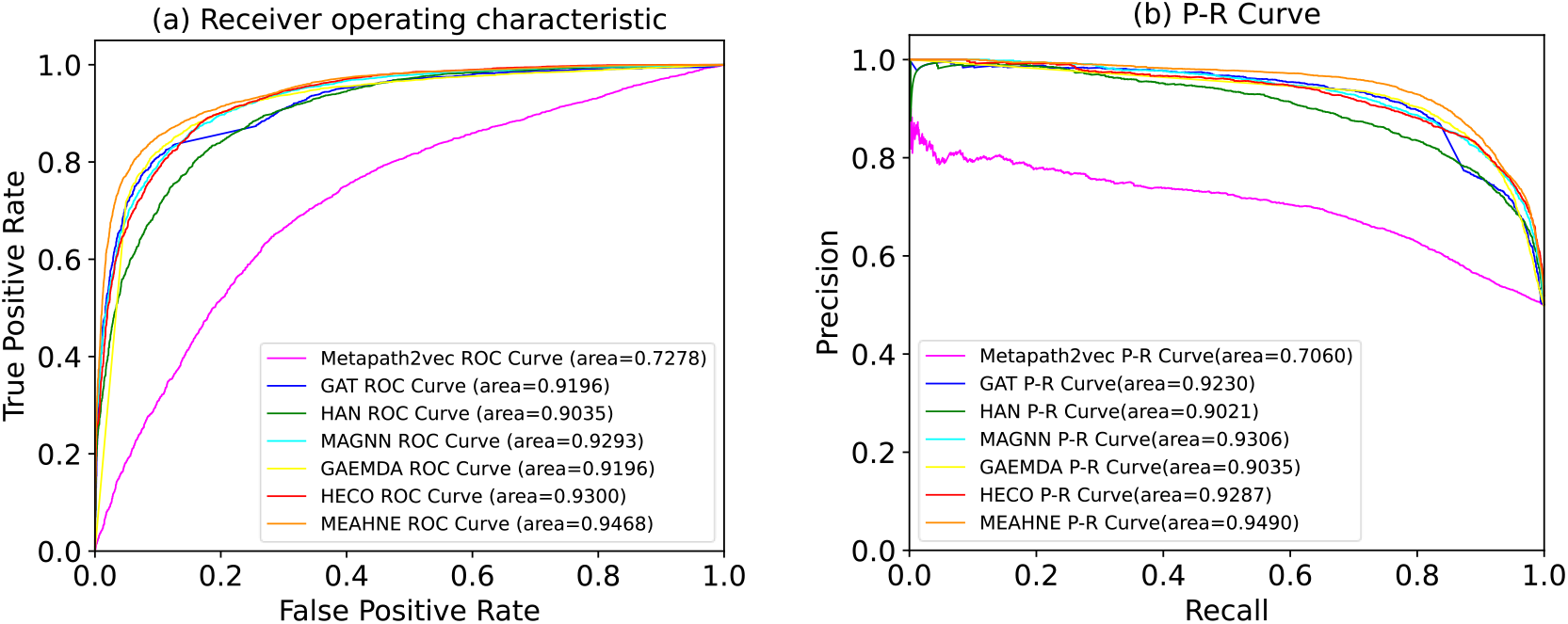
ROC and PR curves for all models **(a)** ROC curves of all models **(b)** P-R curves of all models

### Analysis of experiments

According to the table 1, our model results have good performance under both AUC and AP metrics. We use semantic information twice to make nodes fully aggregated. The first is to encode semantic information into weights to aggregate neighbor nodes, and the second is to fuse semantic information into connected nodes. In this way, our model achieves good performance.

By comparing the experimental results, we can find that since the Metapath2vec model generation node embedding process and the downstream prediction task were performed separately, the downstream prediction task does not affect the generation of upstream nodes. At the same time, the upstream node embedding generation task can only learn the structural information of the node, making the node representation incomplete, which is also the reason why the performance of Metapath2vec was not as good as that of GNN models. The GAT model treats all nodes as being of the same type, which makes GAT unable to learn rich semantic information. It also aggregates all neighbors in the spatial domain, and the noise from the neighbors will also affect the final result. The HAN model only aggregates homogenous nodes connected to the target node through the metapath, which is equivalent to HAN giving semantic information for the meta-path instances and only paying attention to the semantic information at the metapath level. Lack of semantic information leads to the poor performance of HAN. In the GAEMDA model, nodes only aggregate information for connected nodes of different types. Since the auto-encoder continuously updates the node representation on the graph, the node can learn information of nodes that are multiple hops away from it, which helps GAEMDA achieve better results than HAN.

The MAGNN model learns semantic information on the metapath instances and aggregates this information, which enables this model to better learn the complex information in the model; therefore, it yielded good results. The HeCo model learns node representations from two perspectives. Contrastive learning method makes the two perspectives constrain and complement each other in the learning process. The node representation obtained in this way is very complete. HeCo yields good results.

### Analysis of model parameters

We changed two parameters in our model, dimension of the node vector and the number of semantic information extraction layers, to evaluate their influence on performance of the model. In this section, we describe experiments used to evaluate the influence of these two parameters on the model.

For the dimension of the node vector, we found that when the node vector was 64-dimensional, the model performance was better, but when the dimensions were 128 and 256, the model performance became worse (Fig 2). Thirty-two dimensions could not fully express the node information, resulting in loss of information, while 128 and 256 dimensions were too many and contained a lot of noise; 256 dimensions contained the most noise and led to the worst performance.

**Fig 2.**
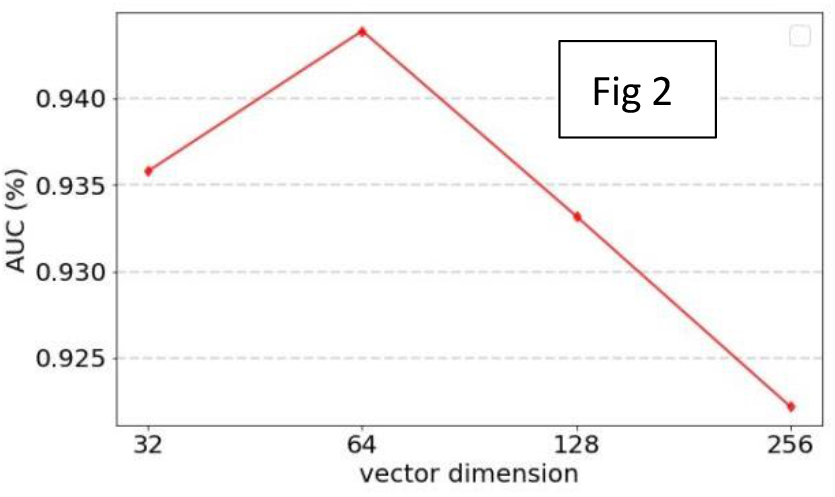
AUC obtained by vector dimention

Our model uses non-linear fully connected layer to build when extracting semantic information on the instance of the metapath, and different number of connection layers affects the quality of information extraction. It can be seen in Fig 3 that the performance was best when the number of node fusion layer was one; the performance of two layers and three layers were similar, and four layers was the worst. Multiple non-linear fully connected layers cause over-fitting, resulting in insufficient semantic information learned. The poor performance of the four layers verifies this point of view.

**Fig 3.**
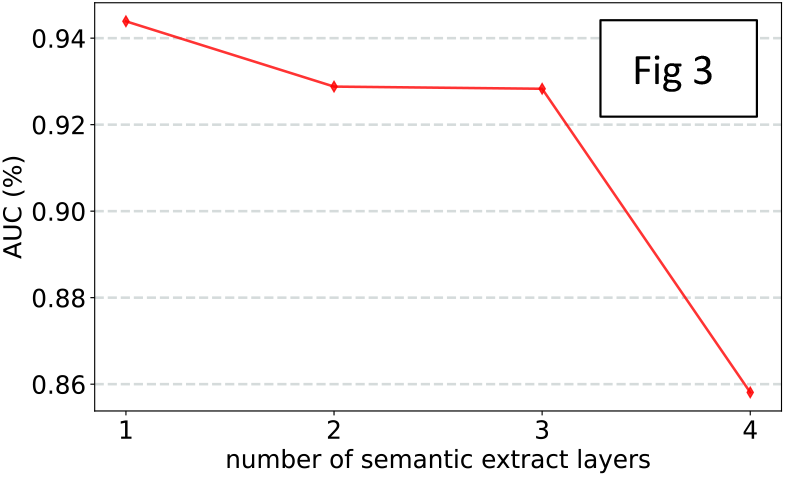
AUC obtained by different number of semantic extract layers

### Case studies

To test the accuracy of our model, we performed miRNA predictions for lung cancer and nasopharyngeal carcinoma. The prediction method was as follows: all of the miRNAs were paired with these two diseases to obtain miRNA-disease pairs, and the trained model was used to score the pairs. We selected the top 50 miRNAs in the miRNA disease combination for evaluation. Among the potential miRNA prediction results for lung cancer, after dbDEMC [34] verification, there were a total of 49 associations with lung cancer. The association of hsa-mir-1-1 was not verified using dbDEMC. Among the potential miRNAs associated with throat cancer, 44 miRNAs in the top 50 miRNAs were verified using the dbDEMC database. The miRNAs that were not verified were hsa-mir-210, hsa-mir-92-1, hsa-mir-1-1, hsa-mir-9-3, hsa-mir-9-2, and hsa-mir-9-1. The prediction results are shown in Tables 2 and 3.

**Table 2.**
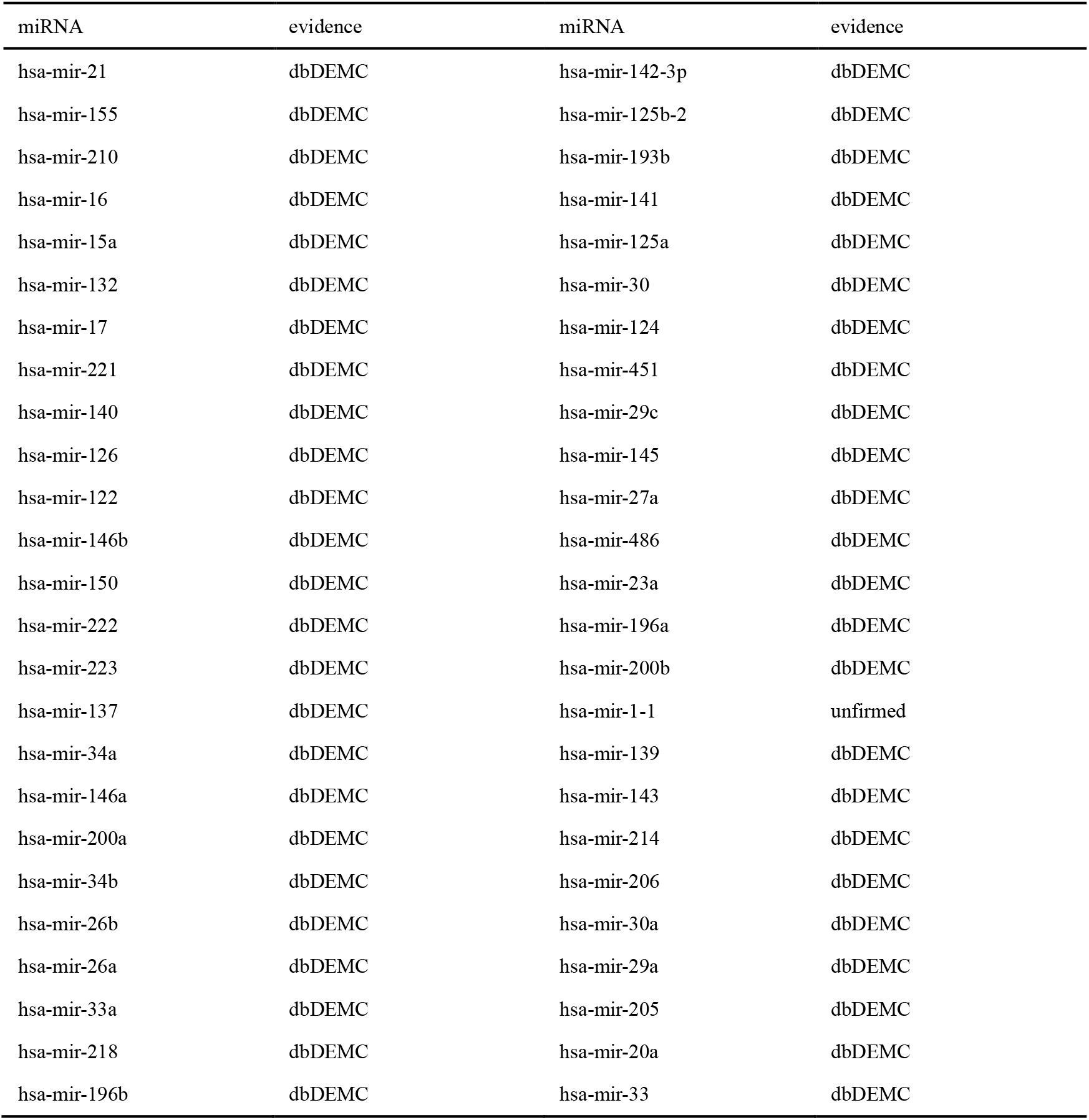
The top 50 miRNAs associated with lung cancer in the prediction results

| miRNA | evidence | miRNA | evidence |
| --- | --- | --- | --- |
| hsa-mir-21 | dbDEMC | hsa-mir-142-3p | dbDEMC |
| hsa-mir-155 | dbDEMC | hsa-mir-125b-2 | dbDEMC |
| hsa-mir-210 | dbDEMC | hsa-mir-193b | dbDEMC |
| hsa-mir-16 | dbDEMC | hsa-mir-141 | dbDEMC |
| hsa-mir-15a | dbDEMC | hsa-mir-125a | dbDEMC |
| hsa-mir-132 | dbDEMC | hsa-mir-30 | dbDEMC |
| hsa-mir-17 | dbDEMC | hsa-mir-124 | dbDEMC |
| hsa-mir-221 | dbDEMC | hsa-mir-451 | dbDEMC |
| hsa-mir-140 | dbDEMC | hsa-mir-29c | dbDEMC |
| hsa-mir-126 | dbDEMC | hsa-mir-145 | dbDEMC |
| hsa-mir-122 | dbDEMC | hsa-mir-27a | dbDEMC |
| hsa-mir-146b | dbDEMC | hsa-mir-486 | dbDEMC |
| hsa-mir-150 | dbDEMC | hsa-mir-23a | dbDEMC |
| hsa-mir-222 | dbDEMC | hsa-mir-196a | dbDEMC |
| hsa-mir-223 | dbDEMC | hsa-mir-200b | dbDEMC |
| hsa-mir-137 | dbDEMC | hsa-mir-1-1 | unfirmed |
| hsa-mir-34a | dbDEMC | hsa-mir-139 | dbDEMC |
| hsa-mir-146a | dbDEMC | hsa-mir-143 | dbDEMC |
| hsa-mir-200a | dbDEMC | hsa-mir-214 | dbDEMC |
| hsa-mir-34b | dbDEMC | hsa-mir-206 | dbDEMC |
| hsa-mir-26b | dbDEMC | hsa-mir-30a | dbDEMC |
| hsa-mir-26a | dbDEMC | hsa-mir-29a | dbDEMC |
| hsa-mir-33a | dbDEMC | hsa-mir-205 | dbDEMC |
| hsa-mir-218 | dbDEMC | hsa-mir-20a | dbDEMC |
| hsa-mir-196b | dbDEMC | hsa-mir-33 | dbDEMC |

**Table 3.**
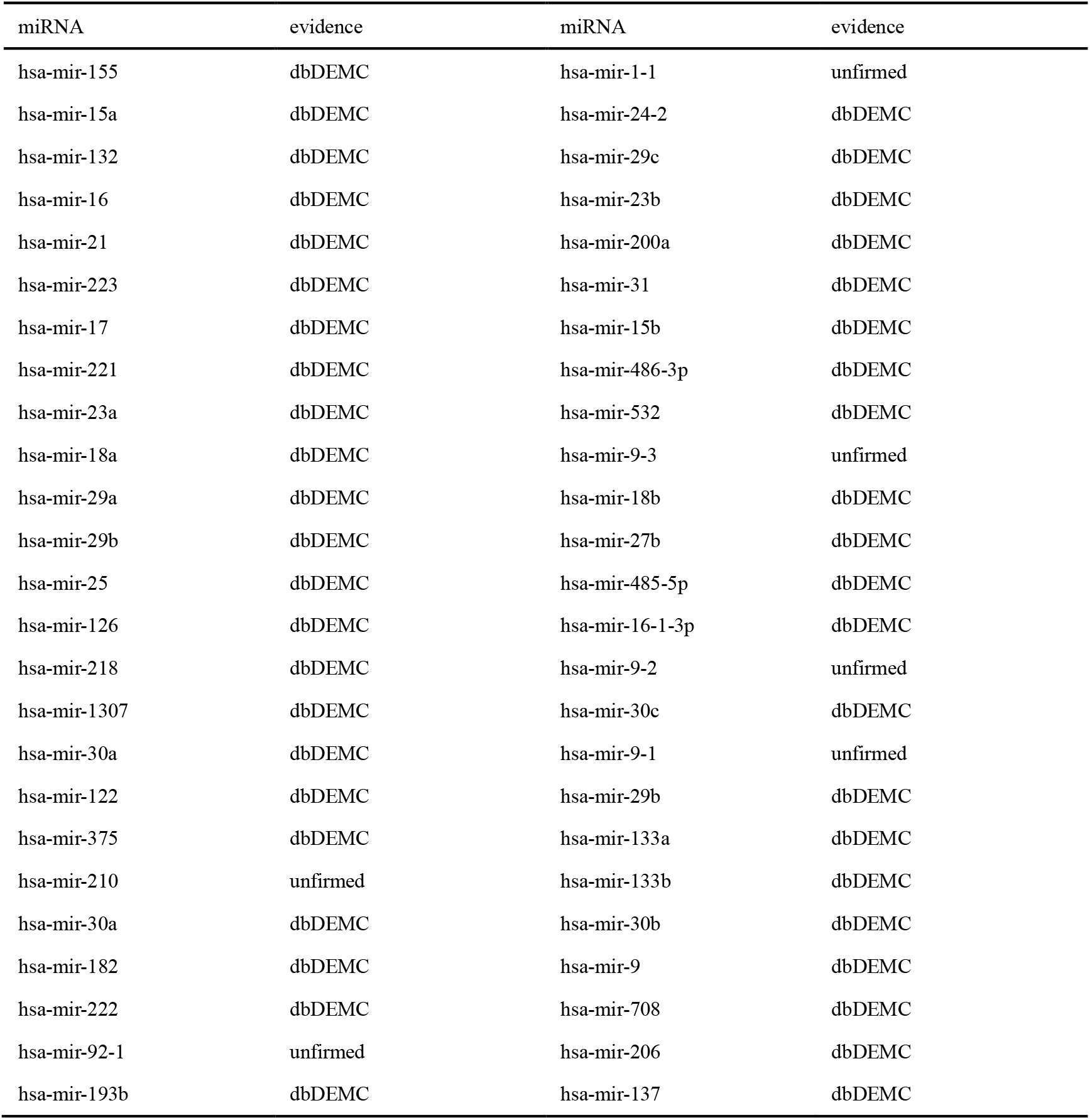
Top 50 miRNAs associated with nasopharyngeal carcinoma in the prediction results

### Conclusion

Due to the relatively small number of verified relationships between miRNA and disease, we selected the third type of node gene and build a heterogeneous network to alleviate this problem.

Meanwhile, we propose a semantic-based heterogeneous graph neural network model for link prediction, which aggregates nodes using semantic-based attention aggregation method. The model utilizes semantic information to aggregate nodes in heterogeneous networks twice, and nodes can fully express the relationships in the network. Traditional heterogeneous graph neural networks often ignore intermediate nodes[25] and cause information loss. We avoid this problem by extracting the information on metapath instances into semantic information. Compared with the heterogeneous graph neural network methods of the past few years, our method achieves the best performance in both AUC and AP.

But it is worth mentioning that our model still has room for improvement. First we randomly select a fixed number of metapath instances for each node. Other selection strategies may yield better performance. Second, semantic information has an important place in our model. Our model uses neural networks to extract semantic information. Whether other extraction methods can make the model perform better is worthy of our further experiments.

### Materials

This section introduces the data we used, which consist of three types of nodes, namely miRNA, disease, and gene, and types kinds of associations between the three types of nodes. The four types of associations are miRNA-disease association, miRNA-gene association, disease-gene association, and protein-protein interaction association.

We collected related links between miRNAs and diseases from the HMDD3.2 [28] database. HMDD is a reliable database that specifically collects miRNA-disease associations. We collected 17,972 links between 1206 miRNA and 893 diseases and integrated miRNAs and diseases as nodes, and miRNA-disease associations as instances into the heterogeneous network. We collected related links between miRNAs and target genes from the Circ2disease [39] database. We selected 4676 links between 202 miRNAs and 1713 genes and integrated miRNAs and target gene as nodes and the associations between them as instances into the heterogeneous network. We collected the related links between diseases and genes from DisGeNET [30]. We selected 84,038 links between 11,181 diseases and 9703 genes and integrated diseases and genes as nodes and the associations between then as instances into the heterogenous network.

The protein-protein interaction network was obtained from the STRING [31] database, which is a reliable database that specifically collects protein interactions. We select genes associated with our chosen miRNAs and diseases and integrate these genes into our heterogeneous network The 105,171 associations between these genes were integrated into the heterogeneous network as instances. Finally, we established a heterogeneous network with 1296 miRNAs, 11,783 diseases, 10,116 genes, and 211,857 instances (Tables 4 and 5).

**Table 4.** Nodes in the network

| Node | Number | Source Dataset |
| --- | --- | --- |
| miRNA | 1296 | HMDD3.2/Circ2disease |
| disease | 11783 | DisGeNET/HMDD3.2 |
| gene | 10116 | Circ2disease/DisGeNET |

**Table 5.**
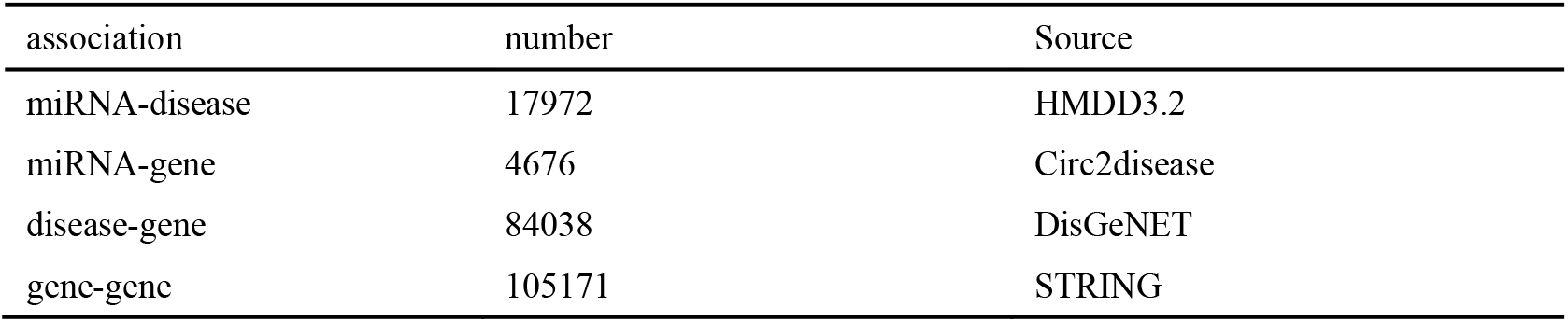
Associations in the network

| association | number | Source |
| --- | --- | --- |
| miRNA-disease | 17972 | HMDD3.2 |
| miRNA-gene | 4676 | Circ2disease |
| disease-gene | 84038 | DisGeNET |
| gene-gene | 105171 | STRING |

### Methods

Definition of metapath. Heterogeneous networks have many types of nodes and many types of relationships. The paths composed of different types of nodes and different types of instances contain rich semantic information, which is not available in homogeneous graphs. To learn the semantic information in heterogeneous networks, the concept of metapath is proposed. For example: 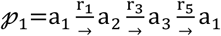 is a metapath, and 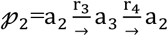 is another metapath, in which 𝓅_i_ (𝓅_i_ ∈ 𝒫) represents a specific metapath, 𝒫 represents all types of metapaths in the heterogeneous network. a_i_ ∈ 𝒜, in which a_i_ represents the i-th type of nodes in the heterogeneous network and 𝒜 represents the collection of all node types in the heterogeneous network. r_i_ ∈ ℛ, r_i_ represents the i-th type of relationship between nodes and ℛ represents the collection of all relationship types in the heterogeneous network.

Definition of metapath instance [26]. Under each metapath type 𝓅_i_, there are a large number of paths following 𝓅_i_ in the heterogeneous network. We call these paths metapath instances. For example, 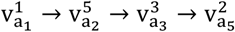 is a metapath instance under 𝓅_1_, in which 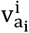 represents the i-th node of type a_i_.

Definition of metapath neighbors. The two endpoints of a metapath instance are metapath neighbors to each other, and they are connected by the metapath instance. For example, 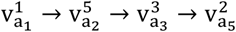 is a metapath instance in which 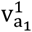 and 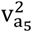 are metapath neighbor to each other.

This section introduces the main methods, ideas, and specific implementation details of the MEAHNE model. The MEAHNE model is mainly divided into five parts: node conversion, subgraph extraction, metapath instance semantic information extraction, node aggregation method based on metapath instance semantic attention, multi-semantic information fusion, and link prediction. Fig 4 shows the overall framework of MEAHNE.

**Fig 4.**
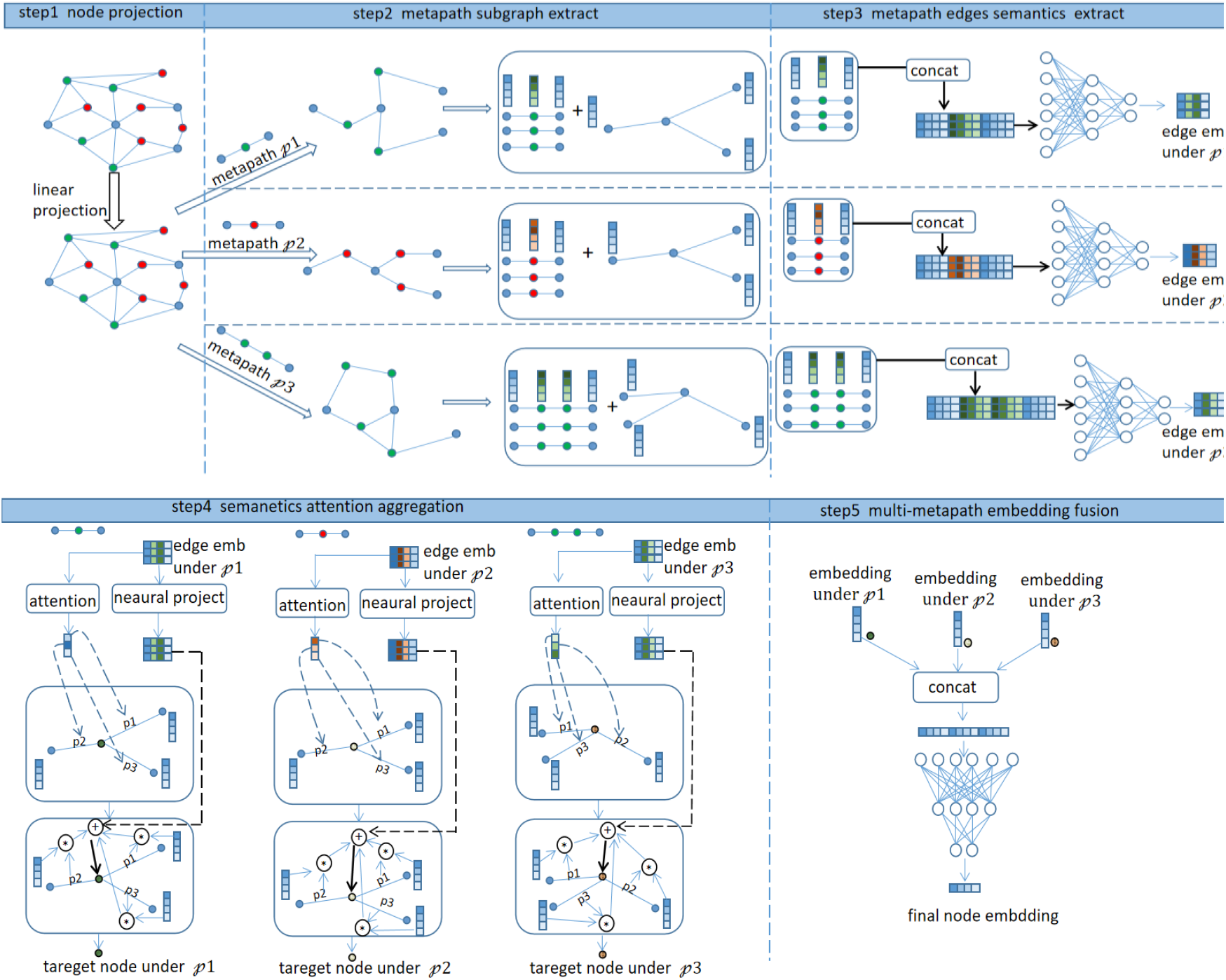
MEAHNE framework. First, nodes of different types are projected into the same space. We use neural networks to extract semantic information in metapath instances. We encoded the semantic information into values as weights to aggregate nodes. The semantic information on metapath instances was aggregated to obtain a more powerful node representation. Finally, representations under all metapaths were fused to obtain the final node embedding.

#### A. Node space conversion

If we want to learn representations of heterogeneous networks, we need to perform interactive calculations on the nodes of the graph. However, heterogeneous network have multiple types of nodes, and different types of nodes are located in different spaces. If the nodes are not processed, the interactive calculation between nodes becomes too difficult, so we first converted all types of nodes into the same space to facilitate calculations between nodes as follows.

A trainable linear transformation matrix was set for each type of node, and original nodes of different types were projected into the same space, as shown in formula (1):

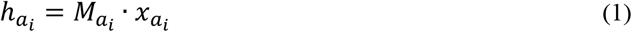

Where 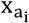 represents the original feature vector of the node type a_i_, and 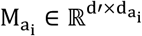, in which d′ represents the feature space dimension after space conversion and 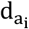 represents the original feature dimension of a_i_ type node.

#### B. Extract the metapath subgraph and the metapath instances

To mine heterogeneous network in multiple metapaths, the first step is to separate the corresponding sub-networks based on specific metapaths.

We separated the sub-network 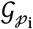 according to the metapath 𝓅_i_, and 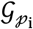 represents the sub-network mined in the 𝓅_i_ mode. In sub-network 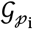, the metapth instances corresponding to the 𝓅_i_ was sampled and denoted as p(v, u), which connects the target node v and its metapath neighbor u.

#### C. Extract the semantic information contained in the metapath instances

When mining the information from the corresponding subgraph 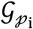 under a single metapath, 𝓅_i_, different types of nodes are transformed into the same space through space, which allows different types of nodes to represent each other. The metapath instance is composed of different types of nodes connected to each other and contains rich semantic information. Therefore, to learn the semantic information on the metapath instance when learning the subgraph, we first integrated the information on the metapath instance. Each metapath instance was represented as a vector that represents the semantic information on the instance. All the nodes on the metapath instance were concatenated according to the order of the metapath, as shown in formula (2):

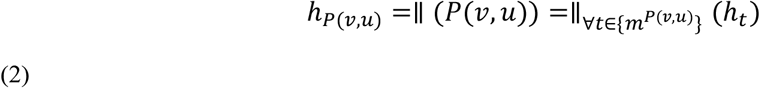

in which *m*^*P*(*ν,u*)^ represents the set of nodes on the metapath instance and *P*(*ν, u*), *h*_*P*(*ν,u*)_ represents the vector obtained by concatenating the vectors of the nodes on the metapath instance *P*(*ν, u*).

The nonlinear neural network was used to learn the vector *h*, and *h* represents the semantic information of the metapath instance. The nonlinear neural network, which has strong information extraction capabilities, is a network composed of multiple fully connected layers and nonlinear activation functions, as shown in formula (3):

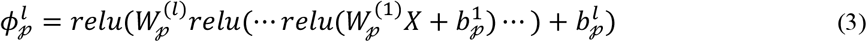

Where 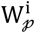 represents the weight matrix of the ith fully connected layer of the neural network under metapath 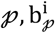 represents the bias value of the ith layer of the neural network under metapath 𝓅, *X* represents the input feature, and 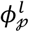 represents the vector representation of the input vector *X* learned through *l* connection layers in the neural network under metapath 𝓅. We used the vector *h*_*P*(*ν,u*)_ as the input of the nonlinear neural network to get the semantic information of metapath instance, as shown in formula (4):

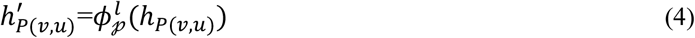

#### D. Semantic-based attention aggregation method

After obtaining the semantic information from the metapath instances, we can aggregate the semantic information into the target nodes connected to these metapath instances, but the semantic information is obtained by the fusion of different types of nodes. If the target node only aggregates semantic information, each type of node contains information about other types of nodes, causing different types of nodes to lose their distinction. To maintain the discrimination between nodes of different types, we first aggregate only neighbor nodes of the same type. For aggregating nodes of the same type, we designed a method to encode semantic information into attention weights and used the obtained attention coefficient to aggregate metapath neighbors. Then, we fused the information obtained by the aggregation of nodes of the same type and semantic information from metapath instances as the final node representation.

We encoded the semantic information on the metapath instance using the attention mechanism as a weight value—the correlation strength coefficient between the target node and the metapath neighbor, as shown in Fig 5 and Equations (5) and (6).

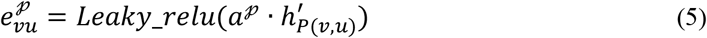

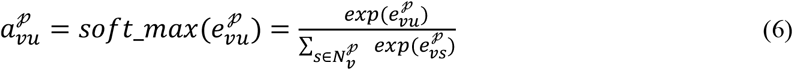

**Fig 5.**
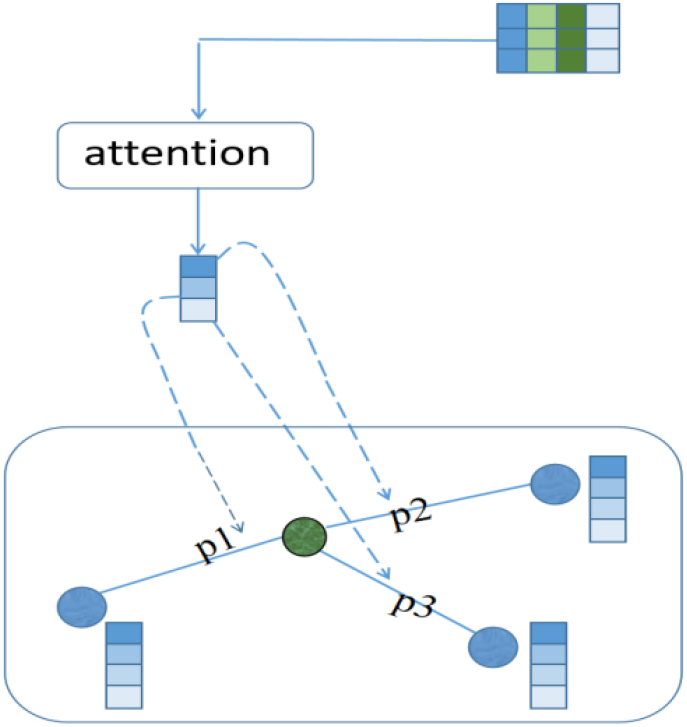
encoding semantic information on the metapth instances into attention weights

Where 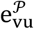 represents the value encoded by the attention mechanism, Leaky_relu (·)is a nonlinear activation function, a^𝓅^ represents the attention weight matrix under metapath 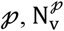 represents the set of metapath neighbors connected to the target node v on the subgraph under metapath 𝓅, and 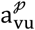 represents the weight value obtained by normalizing 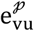.

Next, the metapath neighbors were aggregated according to the weight *a* and the semantic information was also integrated to ensure the integrity of the node embedding.

To reasonably integrate semantic information during the node aggregation stage, we performed secondary learning on semantic information. We designed a trainable matrix to optimize semantic information and added nonlinear activation operations to the optimization results as shown in formula (7).

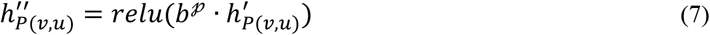

Where *b*^𝓅^ represents a learnable weight matrix under metapath 𝓅, and the content of semantic information is continuously adjusted through end-to-end learning.

Next, the node information was aggregated. We used the learned metapath semantic weight to aggregate the metapath neighbors and added the semantic information learned twice, as shown in formula (8):

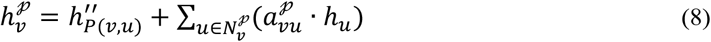

#### E. Fuse embedding of multiple metapaths

In the above steps, we only learned the heterogeneous network under a single metapath. In fact, our model learns the heterogeneous network in multiple metapath modes and generates the representation of the target node in multiple metapath modes. We used neural network methods to integrate node representations under multiple metapaths, as shown by formula (9):

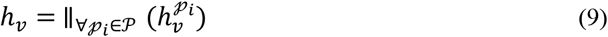

Where 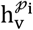 represents the embedding obtained by aggregating the target node v under metapath 𝓅_i_ and*h*_*ν*_ represents the result of concatenating the representation of the target node v under all metapaths. Then the embedding *h*_*ν*_ was input into the nonlinear neural network to learn a low-dimensional embedding that fuses the target node representation under multiple metapaths as shown in formula (10):

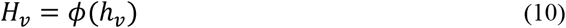

After learning using a nonlinear neural network, *H*_*ν*_ represents a low-dimensional embedding that fuses multiple metapath representation results as the final representation of the target node.

#### F. Link prediction and optimization goals

The vector inner product is used as the score of the link strength of the two nodes. If the two vectors are highly correlated, then the score of the node inner product will be higher. We used this as the basis for link prediction as shown in formula (11):

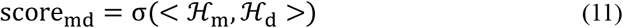

Our link prediction was between miRNA and disease. The higher the prediction score, the stronger the correlation, and the lower the prediction score, the weaker the correlation. In theory this is a two-classification problem, so we used two-class cross-entropy as the optimization target. Our optimization goal is shown in formula (12):

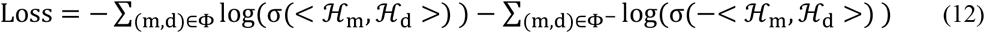

Where Φ represents the set of miRNA and disease pairs that have been verified to be associated, and Φ^-^ represents the set of all miRNA-disease pairs that have not been experimentally verified. The goal of the optimization is to make the score between verified node pair higher and the unverified node pair lower. Because our model is an end-to-end training model, the parameters in the model are continuously optimized during the training process, and the continuously optimized parameters enable us to achieve the optimization goal.

## Competing interests

The authors declare that they have no competing interests.

## Additional Files

All additional files are available at: https://github.com/yyx-hc/MEAHNE

## Authors’ contributions

CH and JL designed the study, performed bioinformatics analysis, and drafted the manuscript. All of the authors performed the analysis. JL conceived of the study, participated in its design and coordination, and drafted the manuscript.

## Acknowlinstancements

This work was supported by the grants from the National Key R&D Program of China (2021YFA0910700), Shenzhen science and technology university stable support program (GXWD20201230155427003-20200821222112001), Shenzhen Science and Technology Program (JCYJ20200109113201726), Guangdong Basic and Applied Basic Research Foundation (2021A1515012461 and 2021A1515220115).

## Key Points

- An attention aggregation method based on metapath instances in heterogeneous networks was developed for miRNA-disease association prediction.
- This model innovatively encodes the semantic information on the metapath instance into weights and uses the obtained weights as the attention for aggregating nodes.
- Heterogeneous neural networks often ignore semantic information on metapath instances. This model fuses the information on the instances to the nodes, which gives the nodes in the network more comprehensive information, and finally calculates the similarity of the nodes in the network to predict the association between miRNAs and disease.

## Notes

### Competing Interest Statement

The authors have declared no competing interest.

## REFERENCES

1. Ambros V. The functions of animal microRNAs[J]. Nature, 2004, 431(7006): 350–355.

2. Lee R C, Feinbaum R L, Ambros V. The C. elegans heterochronic gene lin-4 encodes small RNAs with antisense complementarity to lin-14[J]. cell, 1993, 75(5): 843–854.

3. Lee R C, Ambros V. An extensive class of small RNAs in Caenorhabditis elegans[J]. science, 2001, 294(5543): 862–864.

4. Guo C, Sah J F, Beard L, et al. The noncoding RNA, miR-126, suppresses the growth of neoplastic cells by targeting phosphatidylinositol 3-kinase signaling and is frequently lost in colon cancers[J]. Genes, Chromosomes and Cancer, 2008, 47(11): 939–946.

5. Calin G A, Croce C M. MicroRNA signatures in human cancers[J]. Nature reviews cancer, 2006, 6(11): 857–866.

6. Cahill S, Smyth P, Denning K, et al. Effect of BRAF V600E mutation on transcription and post-transcriptional regulation in a papillary thyroid carcinoma model[J]. Molecular cancer, 2007, 6(1): 1–10.

7. He L, Hannon G J. MicroRNAs: small RNAs with a big role in gene regulation[J]. Nature reviews genetics, 2004, 5(7): 522–531.

8. Goh J N, Loo S Y, Datta A, et al. microRNAs in breast cancer: regulatory roles governing the hallmarks of cancer[J]. Biological Reviews, 2016, 91(2): 409–428.

9. Zhang X, Zeng X. Integrative approaches for predicting microRNA function and prioritizing disease-related microRNA using biological interaction networks[J]. Bio-inspired Computing Models And Algorithms, 2019: 75-105.

10. Chen X, Xie D, Zhao Q, et al. MicroRNAs and complex diseases: from experimental results to computational models[J]. Briefings in bioinformatics, 2019, 20(2): 515–539.

11. Jiang, Y., Liu, B., Yu, L. et al. Predict MiRNA-Disease Association with Collaborative Filtering. Neuroinform 16, 363–372 (2018).

12. Chen X, Yan C C, Zhang X, et al. WBSMDA: within and between score for miRNA-disease association prediction[J]. Scientific reports, 2016, 6(1): 1–9.

13. Chen X, Liu M X, Yan G Y. RWRMDA: predicting novel human microRNA–disease associations[J]. Molecular BioSystems, 2012, 8(10): 2792–2798.

14. You Z H, Huang Z A, Zhu Z, et al. PBMDA: A novel and effective path-based computational model for miRNA-disease association prediction[J]. PLoS computational biology, 2017, 13(3): e1005455.

15. Wu T R, Yin M M, Jiao C N, et al. MCCMF: collaborative matrix factorization based on matrix completion for predicting miRNA-disease associations[J]. BMC bioinformatics, 2020, 21(1): 1–22.

16. Chen X, Wang L, Qu J, et al. Predicting miRNA–disease association based on inductive matrix completion[J]. Bioinformatics, 2018, 34(24): 4256–4265.

17. Kipf T N, Welling M. Semi-supervised classification with graph convolutional networks[J]. arXiv preprint 1609.02907, 2016.

18. Veličković P, Cucurull G, Casanova A, et al. Graph Attention Networks[C]//International Conference on Learning Representations. 2018.

19. Zhang J, Shi X, Xie J, et al. GaAN: Gated Attention Networks for Learning on Large and Spatiotemporal Graphs[C]//34th Conference on Uncertainty in Artificial Intelligence 2018, UAI 2018. 2018.

20. Hamilton W L, Ying R, Leskovec J. Inductive representation learning on large graphs[C]//Proceedings of the 31st International Conference on Neural Information Processing Systems. 2017: 1025–1035.

21. Li J, Zhang S, Liu T, et al. Neural inductive matrix completion with graph convolutional networks for miRNA-disease association prediction[J]. Bioinformatics, 2020, 36(8): 2538–2546.

22. Li Z, Li J, Nie R, et al. A graph auto-encoder model for miRNA-disease associations prediction[J]. Briefings in Bioinformatics, 2021, 22(4).

23. Dong Y, Chawla N V, Swami A. metapath2vec: Scalable representation learning for heterogeneous networks[C]//Proceedings of the 23rd ACM SIGKDD international conference on knowlinstance discovery and data mining. 2017: 135–144.

24. Wang X, Ji H, Shi C, et al. Heterogeneous graph attention network[C]//The World Wide Web Conference. 2019: 2022–2032.

25. Qu Y, Bai T, Zhang W, et al. An end-to-end neighborhood-based interaction model for knowlinstance-enhanced recommendation[C]//Proceedings of the 1st International Workshop on Deep Learning Practice for High-Dimensional Sparse Data. 2019: 1–9.

26. Fu X, Zhang J, Meng Z, et al. Magnn: Metapath aggregated graph neural network for heterogeneous graph embedding[C]//Proceedings of The Web Conference 2020. 2020: 2331–2341.

27. Mikolov T, Chen K, Corrado G, et al. Efficient estimation of word representations in vector space[J]. arXiv preprint 1301.3781, 2013.

28. Huang Z, Shi J, Gao Y, et al. HMDD v3. 0: a database for experimentally supported human microRNA–disease associations[J]. Nucleic acids research, 2019, 47(D1): D1013–D1017.

29. Yao D, Zhang L, Zheng M, et al. Circ2Disease: a manually curated database of experimentally validated circRNAs in human disease[J]. Scientific reports, 2018, 8(1): 1–6.

30. Piñero J, BravoÀ Queralt-Rosinach N, et al. DisGeNET: a comprehensive platform integrating information on human disease-associated genes and variants[J]. Nucleic acids research, 2016: gkw943.

31. Szklarczyk D, Gable A L, Lyon D, et al. STRING v11: protein–protein association networks with increased coverage, supporting functional discovery in genome-wide experimental datasets[J]. Nucleic acids research, 2019, 47(D1): D607–D613.

32. Wang X, Liu N, Han H, et al. Self-supervised Heterogeneous Graph Neural Network with Co-contrastive Learning[J]. arXiv preprint 2105.09111, 2021.

33. Li Z, Li J, Nie R, et al. A graph auto-encoder model for miRNA-disease associations prediction[J]. Briefings in Bioinformatics, 2021, 22(4).

34. Yang Z, Wu L, Wang A, et al. dbDEMC 2.0: updated database of differentially expressed miRNAs in human cancers[J]. Nucleic acids research, 2017, 45(D1): D812–D818.

